# Programmable bacteria induce durable tumor regression and systemic antitumor immunity

**DOI:** 10.1101/561159

**Authors:** Sreyan Chowdhury, Taylor E. Hinchliffe, Samuel Castro, Courtney Coker, Nicholas Arpaia, Tal Danino

**Affiliations:** Department of Biomedical Engineering, Columbia University, New York, NY 10027, USA; Department of Microbiology & Immunology, Vagelos College of Physicians and Surgeons of Columbia University, New York, NY 10032, USA; Herbert Irving Comprehensive Cancer Center, Columbia University, New York, NY 10032, USA; Data Science Institute, Columbia University, New York, NY 10027, USA

## Abstract

Synthetic biology is driving a new era of medicine through the genetic programming of living cells^1,2^. This transformative approach allows for the creation of engineered systems that intelligently sense and respond to diverse environments, ultimately adding specificity and efficacy that extends beyond the capabilities of molecular-based therapeutics^3–5^. One particular focus area has been the engineering of bacteria as therapeutic delivery systems to selectively release therapeutic payloads *in vivo^6–8^*. Here, we engineered a non-pathogenic *E. coli* to specifically lyse within the tumor microenvironment and release an encoded nanobody antagonist of CD47 (CD47nb)^9^, an anti-phagocytic receptor commonly overexpressed in several human cancers^10,11^. We show that intratumoral delivery of CD47nb by tumor-colonizing bacteria increases activation of tumor-infiltrating T cells, stimulates rapid tumor regression, prevents metastasis, and leads to long-term survival in a syngeneic tumor model. Moreover, we report that local injection of CD47nb bacteria stimulates systemic antitumor immune responses that reduce the growth of untreated tumors – providing, to the best of our knowledge, the first demonstration of an abscopal effect induced by a bacteria cancer therapy. Thus, engineered bacteria may be used for safe and local delivery of immunotherapeutic payloads leading to systemic antitumor immunity.

---

The origins of cancer immunotherapy trace back to the pioneering work of Dr. William Coley, who observed tumor clearance in some patients that received injections of bacteria^12^ – a result now attributed to leukocyte activation^13–15^. Since then, a multitude of studies have demonstrated that bacteria preferentially grow within tumor cores due to the immunoprivileged nature of the often hypoxic and necrotic tumor microenvironment^16–18^, and can locally affect tumor growth through the recruitment and activation of leukocytes. This selective colonization has provided an opportunity for using bacteria as drug delivery vehicles for the local expression of recombinant therapeutics in tumors^6^. With the advent of synthetic biology over the past two decades and the development of numerous bacteria gene circuits^19–23^, we reasoned that programming bacteria to controllably release recombinant immunotherapies could allow for local delivery of higher effective concentrations of therapy while preventing toxicities observed following systemic delivery of identical or similar therapeutic agents.

To test the efficacy of this approach, we chose to target CD47, a potent anti-phagocytic receptor overexpressed in several human cancers^10,11,24^. Recent studies have shown that CD47 blockade not only increases phagocytosis of cancer cells but also promotes cross presentation of tumor antigens by dendritic cells to enhance priming of antitumor effector T cells in syngeneic murine tumor models^25–27^. However, as demonstrated in both preclinical models^11^ and human trials^28,29^, CD47 blockade using systemically delivered antibodies can result in anemia and thrombocytopenia due to high expression of CD47 on red blood cells and platelets respectively. To improve upon its therapeutic profile, a nanobody (camelid single heavy chain antibody fragment) against CD47 with ~200-fold higher binding affinity than the commercially available antimouse CD47 monoclonal antibody (miap301) was recently developed and characterized^9^. This nanobody demonstrated mild effects as a systemically administered monotherapy, potentially due to lack of Fc-mediated effector function^30^; however, a notable therapeutic response was observed when used in combination with a tumor specific antibody and systemic immune checkpoint blockade. In this work, we engineered an *E. coli* strain containing a synchronized lysis circuit (eSLC) that colonizes tumors and undergoes intratumoral quorum-lysis to locally release an encoded nanobody antagonist of CD47 (eSLC-CD47nb) (**Fig. 1a**). This system allows for the combined local delivery of a novel immunotherapeutic along with immunostimulatory bacterial lysis adjuvants to stimulate antitumor immunity and promote tumor regression.

**Figure 1.**
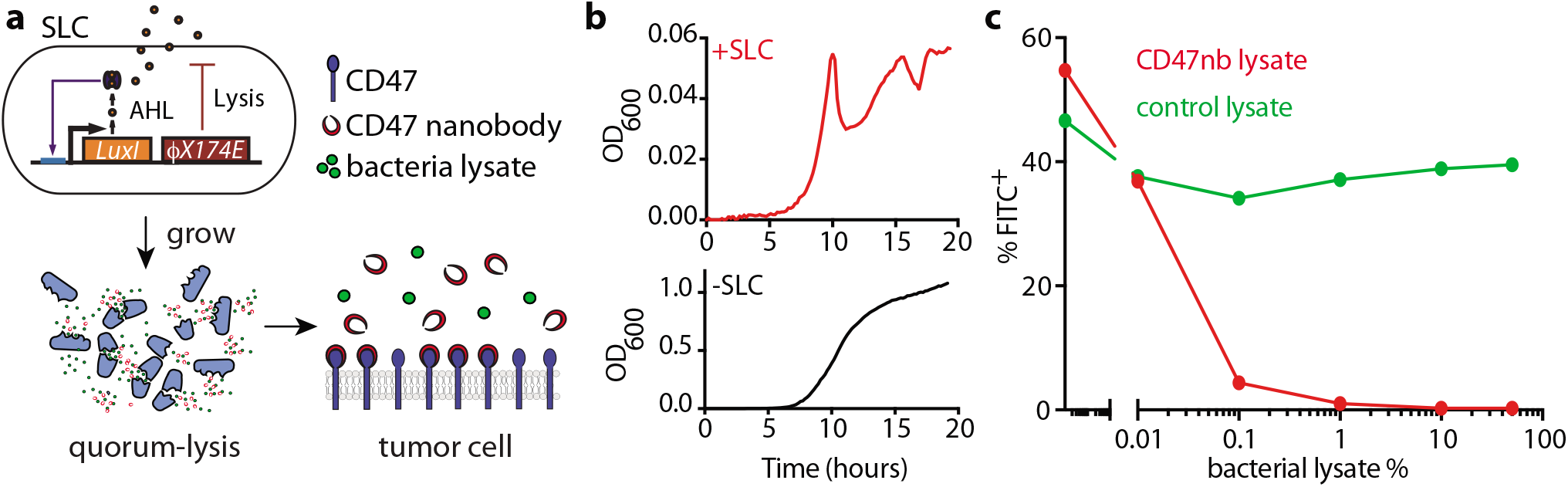
Quorum-induced release of functional anti-CD47 blocking nanobody by engineered immunotherapeutic bacteria encoding a synchronized lysis circuit (SLC). **a**, *E. coli* with SLC reach a quorum and induce the phage lysis protein ϕX174E, leading to bacterial lysis and release of constitutively produced, anti-CD47 blocking nanobody which binds to CD47 on the tumor cell surface. **b**, Bacterial growth dynamics over time of +SLC and -SLC *E. coli* in batch liquid culture. **c**, A20 cells were co-incubated with constant concentration of FITC conjugated αCD47 monoclonal antibody (FITC-miap301) along with varying concentrations of bacterial lysates containing constitutively expressed CD47nb (pSC02) or empty vector (pSC03).

To confirm expression and lysis-dependent release of CD47nb, we first transformed non-pathogenic *E. coli* with a single plasmid encoding the synchronized lysis circuit (eSLC), as well as a stabilized plasmid driving constitutive expression of a hemagglutinin (HA)-tagged variant of CD47nb (**Supplementary Fig. 1**). The SLC strain grows and produces the quorum-sensing molecule acylhomoserine lactone (AHL) via expression of *luxI*, then lyses at a critical threshold due to the production of a bacteriophage lysis protein (*ϕ*x174*E*), resulting in bacterial death and therapeutic release^7^ (**Fig. 1a**). Since this gene circuit was previously tested in *S. typhimurium* on two plasmids of differing copy numbers^7^, we first assessed SLC-mediated lysis of eSLC-CD47nb *E.coli* by time-lapse microscopy of bacteria using an agar pad to limit cell motility^31^. eSLC-CD47nb grew, reached quorum and lysed over a 20-hour time course, in contrast to non-SLC *E. coli* (-SLC) which continuously grew and filled the field of view (**Supplementary Fig. 2a**). Furthermore, we cultured +SLC and -SLC *E. coli* in LB broth in a 96-well plate and measured optical density (OD_600_) over time. eSLC-CD47nb (+SLC) exhibited multiple periodic dips in OD_600_, indicating rounds of synchronized lysis, whereas -SLC *E. coli* exhibited normal bacterial growth kinetics (**Fig. 1b**). Upon verifying synchronized lysis behavior, we evaluated the lysis-mediated release of CD47nb in batch cultures. Immunoblots of log-phase bacterial cultures indicated that eSLC-CD47nb bacteria released significantly higher levels of HA-tagged CD47nb into culture supernatants than control CD47nb-HA bacteria without SLC (**Supplementary Fig. 2b**), suggesting that CD47nb release is enhanced by eSLC. To verify that bacterially-produced nanobody functionally binds CD47, A20 murine lymphoma cells, known to express CD47^25^, were incubated with a fixed concentration of FITC-labeled anti-mouse CD47 mAb (clone miap301) and varying dilutions of bacterial lysate from eSLC-CD47nb or eSLC expressing and empty vector (**Fig. 1c** and **Supplementary Fig. 2c**). We observed a progressive reduction in CD47 staining when cells were incubated with lysates from bacteria expressing CD47nb, suggesting that bacterially produced CD47nb could effectively outcompete miap301 binding to CD47 on the surface of A20 cells. Overall, these results indicate that SLC-CD47nb releases CD47nb in a lysis-dependent manner and that CD47nb produced by engineered eSLC-CD47nb is capable of binding a similar CD47 epitope on tumor cells as the therapeutically effective miap301 mAb.

**Figure 2.**
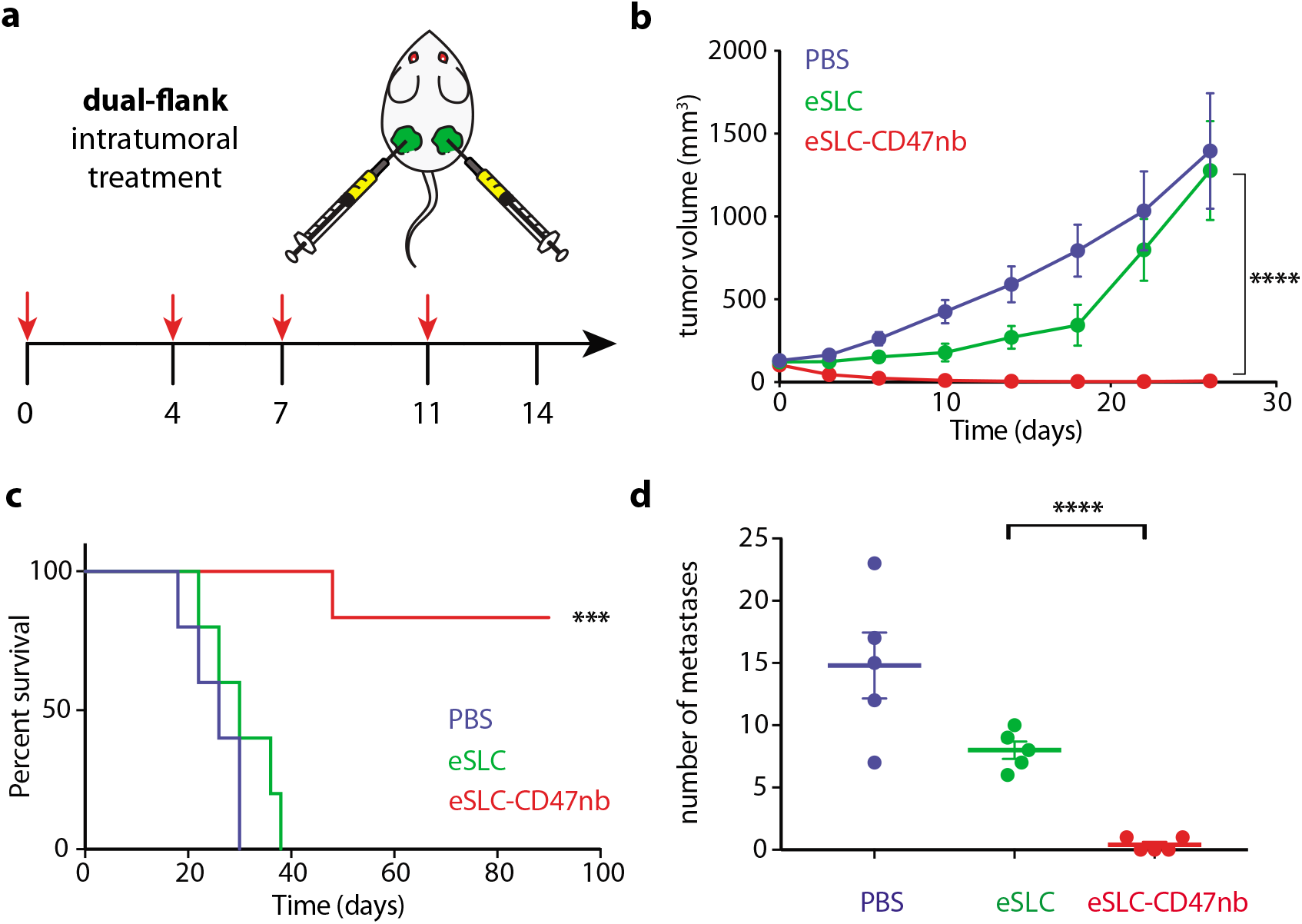
Intratumoral production of CD47 nanobody by eSLC elicits durable tumor regression in a syngeneic murine model of B-cell lymphoma. **a**, Treatment schedule. BALB/c mice (n=7 per group) were implanted subcutaneously with 5 × 10^6^ A20 cells on both hind flanks. When tumor volumes were 100–150 mm^3^, mice received intratumoral injections every 3–4 days with PBS, eSLC or eSLC-CD47nb in both tumors **b**, Tumor growth curves (**** P<0.0001, twoway ANOVA with Tukey’s multiple comparisons test, error bars represent s.e.m.). **c**, Kaplan-Meier survival curves for A20 tumor bearing mice (n=5 per group, ***P<0.001, Log-rank (Mantel-Cox test)). **d**, Quantification of metastatic nodules present in livers on day 30 following bacterial therapy (n= 5 per group, **** P<0.0001, unpaired t-test).

We next sought to evaluate the clinical efficacy for eSLC-CD47nb bacteria in a syngeneic mouse model. BALB/c mice were implanted with 5 × 10^6^ A20 cells in both hind flanks. When tumors reached 100–150 mm^3^ in volume, mice were randomly divided into three groups (PBS, eSLC, eSLC-CD47nb) and received intratumoral injections of PBS, or 10^7^ colony forming units (CFU) of eSLC or eSLC-CD47nb bacteria resuspended in PBS, every 3–4 days for a total of 4 doses (**Fig. 2a**). While administration of control eSLC alone initially slowed tumor growth, likely due to the activation of innate immune cells by bacterial products released upon quorum-lysis, final tumor volumes were not statistically different from PBS treated mice. In contrast, administration of eSLC-CD47nb resulted in rapid and durable clearance of established A20 tumors within ~10 days of commencing therapy (**Fig. 2b** and **Supplementary Fig. 3a and 3b**). Significant therapeutic efficacy was also observed when eSLC-CD47nb was intratumorally injected in a syngeneic murine model of triple negative breast cancer (TNBC) (**Supplementary Fig. 4a–d**). Of note, ~80% of mice treated with eSLC-CD47nb survived >90 days (**Fig. 2c**) and surviving mice were resistant to rechallenge upon subcutaneous injection of 10 × 10^6^ A20 cells (data not shown), while naïve mice receiving the same batch of cells developed tumors within a week of injection. Furthermore, in contrast to animals receiving intratumoral injections of PBS or control eSLC bacteria, liver metastases were rarely observable in mice treated with eSLC-CD47nb when monitored at 30 days post-treatment (**Fig. 2d**). Additionally, lung metastases were notably reduced in mice receiving eSLC-CD47nb in the 4T1 TNBC model (**Supplementary Fig. 4b–d**). We also assessed the biodistribution of eSLC-CD47nb bacteria 72 hours post treatment. Growth remained restricted to tumors (~2 × 10^8^ CFU) and no bacteria could be cultured from the livers or spleens of treated mice above the limit of detection (~1 × 10^3^ CFU) (**Supplementary Fig. 5b**). Furthermore, bacterial therapy was well-tolerated by animals with no changes in body weight observed over the course of treatment or throughout the observation period (**Supplementary Fig. 3b**). Overall, these results indicate that treatment of tumors with eSLC-CD47nb promotes local tumor regression while also preventing metastasis, suggesting the induction of systemic antitumor immunity that is likely to be mediated by tumor-specific T cells.

**Figure 3.**
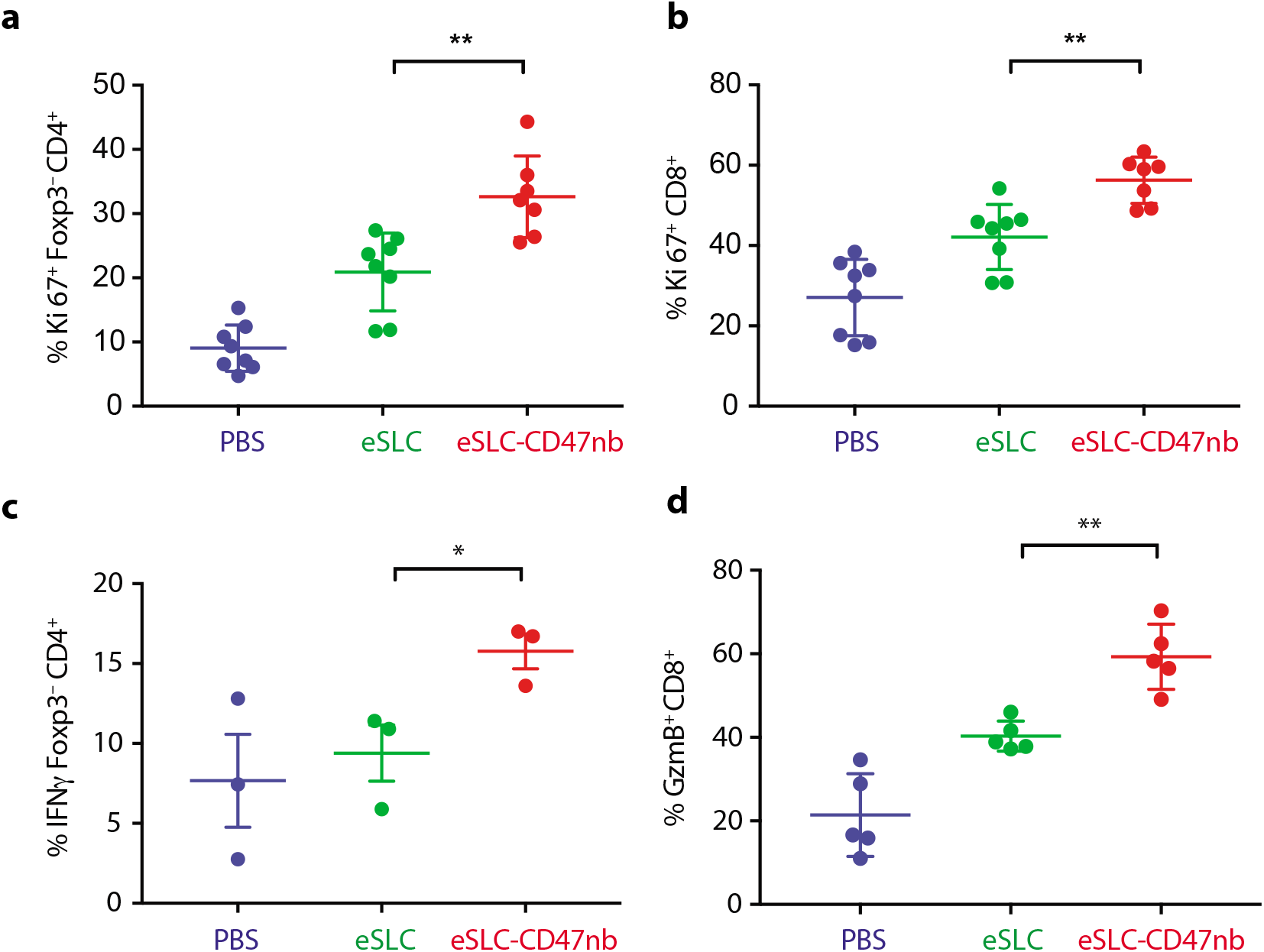
Immunotherapeutic eSLC-CD47nb bacteria prime robust adaptive anti-tumor immune responses. 5 × 10^6^ A20 cells were implanted into the hind flanks of BALB/c mice. When tumors reached 100-150 mm^3^ in volume (day 0), mice were treated with either PBS, eSLC or eSLC-CD47nb on days 0, 4 and 7. On day 8 tumors were homogenized and tumor-infiltrating lymphocytes were isolated for flow cytometric analysis on day 8. **a, b** Frequencies of isolated intratumoral Ki-67^+^ Foxp3^−^CD4^+^ and CD8^+^ T cells. **c**, Tumor infiltrating lymphocytes were stimulated following *ex vivo* isolation with PMA and ionomycin in the presence of brefeldin A. Frequencies of intratumoral IFNγ^+^ Foxp3^−^CD4^+^ T cells following stimulation. **d**, Percentages of intratumoral Granzyme-B positive CD8^+^ T cells. (n= 3-7 per group. * P<0.05, ** P<0.01, unpaired *t*-test).

Recent studies have highlighted the importance of dendritic cells and effector T cells as being indispensable for anti-CD47–mediated clinical responses^25,26^. We reasoned that local inflammation induced by bacterial lysis coupled with localized blockade of CD47 on tumor cells would increase tumor cell phagocytosis and tumor antigen presentation, and thereby enhance the priming of antitumor T cells. Indeed, when we compared the effect of systemically delivered CD47 antibody (miap301) we observed no tumor regression or enhanced survival, whereas mice receiving intravenous eSLC-CD47nb showed significantly slower tumor growth (**Supplementary Fig. 6b**). Moreover, immunophenotyping of A20 tumors treated with eSLC-CD47nb revealed increased proliferation of both Foxp3^−^CD4^+^ and CD8^+^ T cells (**Fig. 3a and 3b**) in comparison to tumor bearing mice treated with eSLC bacteria. Furthermore, tumor-infiltrating Foxp3^−^CD4^+^ T cells from eSLC-CD47nb-treated tumors produced significantly higher levels of IFN-γ following *ex vivo* restimulation with PMA and ionomycin (**Fig. 3b and 3c**). While eSLC-CD47nb did not lead to significant changes in IFN-γ levels in CD8^+^ T cells (**Supplementary Fig. 7f**), we observed markedly elevated levels of intratumoral Granzyme B^+^ CD8^+^ T cells (**Fig. 3d**). Additionally, these T cell responses appeared to be tumor antigen-specific as overnight *in vitro* co-culture of irradiated A20 cells with splenocytes derived from mice treated with eSLC-CD47nb led to robust secretion of IFN-γ (**Supplementary Fig. 8**). Interestingly, mice treated with miap301 exhibited no elevated IFN-γ response in comparison to those receiving eSLC-CD47nb. These data suggest that eSLC-CD47nb not only support the activation (**Supplementary Fig. 7a and 7b**) and proliferation of intratumoral T cells, but also lead to the induction of systemic anti-A20 memory T cell responses.

Durable remission from cancer requires not only elimination of treated tumors but also systemic antitumor immunity for the clearance of distant metastases. Based on our observation that eSLC-CD47nb enhances the effector function of tumor-infiltrating T cells within treated tumors, we examined whether SLC-CD47nb could delay growth of untreated tumors. Mice injected with A20 tumors on both flanks were treated with eSLC-CD47nb in a unilateral fashion (**Fig. 4a**). While control eSLC bacteria had no effect on the growth of untreated tumors, treatment of the primary tumor with eSLC-CD47nb substantially slowed the growth of untreated tumors on the opposing flank (**Fig. 4b and 4c**, **Supplementary Fig. 5a**). To further quantify this effect, we computed the mean growth rates (mm/day) of treated vs. untreated tumors by calculating the slopes in tumor volume trajectories for each mouse (**Fig. 4c**). These data indicated that eSLC-CD47nb treated mice showed decreased growth rates in both treated and untreated tumors compared to controls. We considered the possibly that SLC-CD47nb bacteria injected into the primary tumor migrated via systemic circulation to seed the untreated lesion; however, both at 72 hours and 30 days postinjection, we were unable to detect bacteria within untreated tumors growing on the opposing flank (**Supplementary Fig. 5b**), indicating that an adaptive immune response may be mediating this abscopal effect. In support of this hypothesis, flow cytometric analysis of lymphocytes isolated from untreated tumors of mice whose primary tumors were injected with eSLC-CD47nb showed increased frequencies of activated Ki67^+^CD8^+^ T cells (**Fig. 4d** and **Supplementary Fig. 9d**), with a significantly higher percentage of CD8^+^ T cells producing IFN-γ following *ex vivo* restimulation with PMA and ionomycin (**Fig. 4e**).

**Figure 4.**
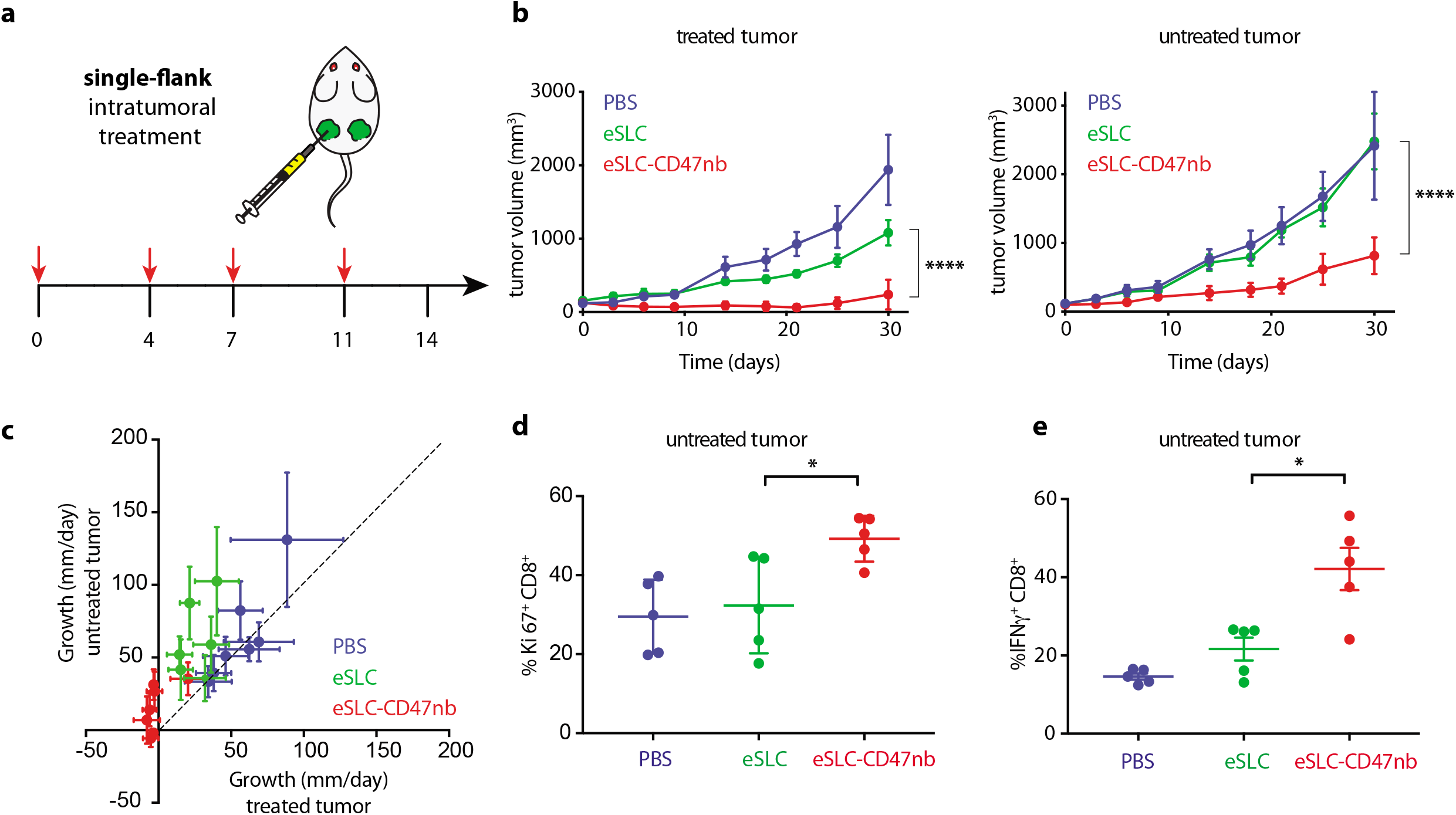
Systemic adaptive immunity following bacterial therapy limits growth of untreated tumors. **a**, treatment schedule. BALB/c mice (n=4 per group) were implanted subcutaneously with 5 × 10^6^ A20 cells on both hind flanks. When tumor volumes reached 100-150 mm^3^, mice received intratumoral injections every 3–4 days with PBS, eSLC or eSLC-CD47 into a single tumor. **b**, tumor growth curves (**** P<0.0001, two-way ANOVA with Tukey’s multiple comparisons test, error bars represent s.e.m.). **c**, Plot of untreated tumor growth rate (mm/day) vs. treated tumor growth rate (mm/day) for each mouse. Dotted line indicates slope=1, points represent means, error bars represent s.e.m. **d**, Untreated tumors were isolated on day 8 following single flank bacterial injections and analyzed by flow cytometry (n=5 per group). Frequencies of intratumoral IFN-γ^+^ Foxp3^−^CD4^+^ and CD8^+^ tumor infiltrating T cells following *ex vivo* stimulation with PMA and ionomycin in the presence of Brefeldin A (* P<0.05, unpaired t-test).

The approach described herein couples the inherently immunostimulatory nature of bacterial lysis products from programmable bacteria with potent nanobody-mediated blockade of an antiphagocytic receptor. Consequently, we observed enhanced infiltration and activation of tumor infiltrating lymphocytes leading to the induction of durable and systemic anti-tumor immunity. Our results suggest that localized, lysis-mediated release of anti-CD47 nanobody confers multiple advantages over conventional systemic monoclonal antibody therapy – firstly, intratumoral delivery of nanobody by eSLC increases the local concentration of immunotherapy while simultaneously preventing systemic toxicity. Second, local treatment with eSLC-CD47nb promotes the induction of systemic antitumor immune responses that are not observed following treatment with anti-CD47 monoclonal antibody. Finally, the ease of engineering bacteria to express additional immunotherapeutic nanobodies and/or cytokines, opens the possibility of evaluating combinations of several other immunotherapeutics which have exhibited systemic toxicity but may be safe and effective when delivered intratumorally using eSLC. The system we describe allows for the delivery of immunotherapeutics in a spatiotemporally defined manner and permits their delivery within diverse solid tumor settings. Moreover, due to the observed abscopal effect, this suggests a future strategy for treating metastatic lesions through the injection of accessible primary tumors.

## METHODS

### Strains and Plasmids

Plasmids were constructed using Gibson assembly or standard restriction enzyme-mediated cloning methods. The pSC01 SLC plasmid was constructed by first amplifying a region containing the constitutively expressed *luxR* gene and phage lysis gene, *ϕx174E* under the control of *luxI* promoter from a pZA35E plasmid^7^. Next, this was cloned into a pTD103-luxI plasmid^22^ using the AvrII site. The pSC02 therapeutic plasmid was constructed by cloning a gBlock (IDT) encoding a *tac* promoter and an *E. coli* codon-optimized sequence for the A4 anti-CD47 nanobody^9^ with an C-terminal hemagglutinin tag into the multiple cloning site of a pAH162 plasmid^32^. Additionally, two stabilizing elements, the hok/sok system^33^ and alp7 partitioning system^34^ were introduced into pSC02 to minimize plasmid loss *in vivo*. pSC01 and pSC02 were transformed into chemically competent *E. coli* Pir1^+^ (Invitrogen). eSLC strains were grown in LB media with 50 μg/mL kanamycin (pSC01) and 100 μg/mL tetracycline (pSC02) along with 0.2% glucose at 37°C for under 12 hours in a shaking incubator. Glucose was added to reduce expression from the Lux promoter and prevent lysis *in vitro*.

### Synchronized Lysis Circuit (SLC) characterization

To validate SLC function, +SLC and -SLC *E. coli* were inoculated into LB media containing appropriate antibiotics and diluted 1:10. Samples were grown at 37°C in a round bottom 96 well plate in a shaking Tecan plate reader. OD_600_ was recorded every 10 minutes for 20 hours. Agar pads were prepared according to previous protocols^31^. +SLC and -SLC *E. coli* were inoculated into LB media containing appropriate antibiotics and grown to mid-log phase. They were diluted 1:100 and grown under agar pads at 37°C and imaged using a Nikon Ti-E microscope equipped with an Okolab stage top incubator.

### Bacterial nanobody characterization

Overnight cultures of *E. coli* containing pSC01 and pSC02 were grown in appropriate antibiotics and 0.2% glucose. A 1:100 dilution into LB with antibiotics was made the following day and bacteria were grown in a shaking incubator at 37 °C. Optical Density (OD) was measured every 30 mins until the OD of lysing strains began to fall, indicating lysis. At this point OD normalized bacteria were spun down at 3000 rcf and supernatants were filtered through a 0.2 μm filter. Cell pellets were mechanically lysed by 3–4 freeze-thaw cycles. Supernatants and lysates were separated by SDS-PAGE followed by immunoblotting with rat anti-HA (Sigma) antibody to evaluate the presence of recombinant CD47nb protein in culture fractions. To verify binding of bacterially produced nanobody to CD47 on tumor cells, serial dilutions of bacterial supernatants from SLC bacteria with or without pSC02 were co-incubated with FITC-labeled anti-CD47 antibody in the presence of A20 tumor cells for 1 hour and FITC fluorescence was measured by flow cytometry.

### Animal models

All animal experiments were approved by the Institutional Animal Care and Use Committee (Columbia University, protocol AC-AAAN8002). The protocol requires animals to be euthanized when tumor burden reaches 2000 mm^3^ or under veterinary staff recommendation. Mice were blindly randomized into various groups. Animal experiments were performed on 4–6 week-old female BALB/c mice (Taconic Biosciences) with bilateral subcutaneous hind flank tumors from A20 murine lymphoma cells (ATCC). The concentration for implantation of the tumor cells was 5 × 10^7^ cells per ml in RPMI (without phenol red). Cells were injected subcutaneously at a volume of 100 μl per flank, with each implant consisting of 5 × 10^6^ cells. Tumors were grown to an average volume of approximately 100–150 mm^3^ before treatment with eSLC strains. Tumor volume was calculated by measuring the length and width of each tumor using calipers, where V = length x width^2^ × 0.5 as previously calculated^25^. The growth in mm/day was computed by taking the difference between tumor volumes at adjacent time points for a particular animal. Values were computed as the mean with standard error plotted.

### Bacterial administration for *in vivo* experiments

Bacterial strains were grown overnight in LB media containing appropriate antibiotics and 0.2% glucose. A 1:100 dilution into media with antibiotics was started the day of injection and grown to an OD_600_ of approximately 0.1. Bacteria were spun down and washed 3 times with sterile PBS before injection into mice. Intratumoral injections of bacteria were performed at a concentration of 5 × 10^8^ CFU per ml in PBS with a total volume of 20–40 μl injected per tumor.

### Flow cytometry

Tumors were extracted for immunophenotyping on day 8 following commencement of bacterial therapy. Lymphocytes were isolated from tumor tissue by mechanical homogenization of tumor tissue followed and digestion with collagenase A (1 mg/ml; Roche) and DNase I (0.5 μg/ml; Roche) in isolation buffer (RPMI 1640 supplemented with 5% FBS, 1% l-glutamine, 1% pen-strep and 10 mM Hepes) for 1 hour at 37°C. Cells were filtered through 100 μm cell strainers, washed in isolation buffer and stained. Dead cells were excluded by staining with Ghost Dye cell viability reagent. Extracellular antibodies used included anti-B220 (BD), anti-CD4 (Tonbo), anti-CD8 (eBioscience) and anti-NKp46 (BD). To measure T cell production of cytokines, cells were stimulated for 2 hours with PMA (50 ng/ml Sigma), ionomycin (1nM; Calbiochem) in the presence of GolgiPlug (brefeldin A). Following extracellular staining with the aforementioned antibodies, intracellular staining was performed using anti-CD3 (Tonbo) anti-TCRβ (BD), anti-CTLA4 (eBioscience), anti-Foxp3 (eBioscience), anti-Ki-67 (Thermo), and cytokines (anti-IL-17 (eBioscience), anti-TNF-α (eBioscience), anti-IFN-γ (Tonbo). Cells were fixed using Foxp3/transcription factor staining buffer set (Tonbo) as per manufacturer’s protocol. Samples were analyzed using a BD LSRFortessa cell analyzer.

### Statistical analysis

Statistical tests were calculated in GraphPad Prism 7.0 (Student’s *t*-test and ANOVA). The details of the statistical tests carried out are indicated in the respective figure legends. Where data were approximately normally distributed, values were compared using either a Student’s *t*-test or oneway ANOVA for single variable, or a two-way ANOVA for two variables with Tukey’s correction for multiple comparisons. For Kaplan-Meier survival experiments we performed a Log-rank (Mantel-Cox) test. Mice were randomized in different groups before experiments.

### Data availability

The data that support the findings of this study are available within the paper and its supplementary information files. Additional data are available from the authors upon reasonable request.

## ACKNOWLEDGEMENTS

This work was supported by the NIH Pathway to Independence Award (R00CA197649-02) (T.D.), DoD Idea Development Award (LC160314) (T.D.), DoD Era of Hope Scholar Award (BC160541) (T.D.), NIH AI127847 (N.A.), Searle Scholars Program SSP-2017-2179 (N.A.), and the Roy and Diana Vagelos Precision Medicine Pilot Grant (N.A. and T.D.). Research reported in this publication was performed in the Columbia University Department of Microbiology & Immunology Flow Cytometry Core facility. The content is solely the responsibility of the authors and does not necessarily represent the official views of the National Institutes of Health. We would like to thank Katherine T. Fortson and Oscar Velasquez for technical assistance with flow cytometry experiments and *in vivo* tumor experiments respectively. We thank Rosa L. Vincent and Thomas M. Savage for review of the manuscript.

## AUTHOR CONTRIBUTIONS

S.C., N.A. and T.D. conceived and designed the study. S.C., T.E.H., C.C., and S.C. performed *in vivo* experiments. S.C performed *in vitro* characterization of eSLC-CD47nb and conducted immunophenotyping experiments. S.C., N.A. and T.D. analyzed data and wrote the manuscript with input from all other authors.

## COMPETING INTERESTS STATEMENT

S.C., N.A. and T.D. have filed a provisional patent application with the US Patent and Trademark Office (US Patent Application No. 62/747,826) related to this work.

## SUPPLEMENTARY FIGURES FOR

**Supplementary Figure 1.**
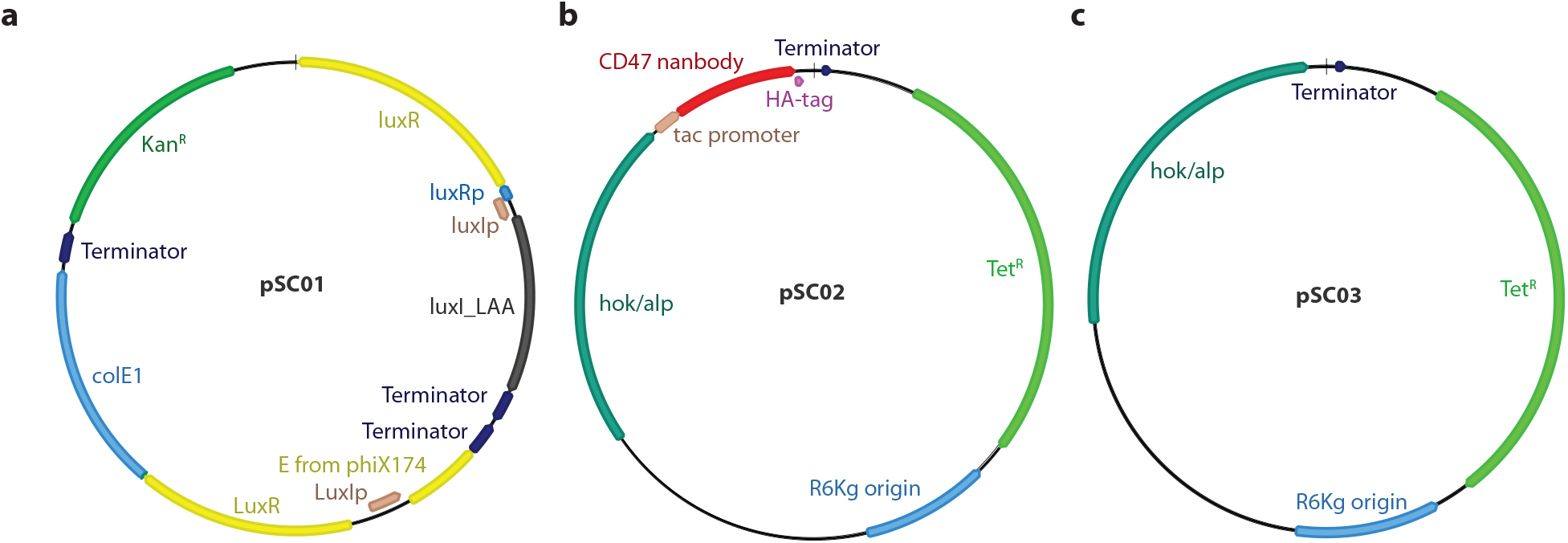
Map of plasmids used in this study. **a**, pSC01, single plasmid synchronized lysis circuit. **b**, pSC02, stabilized plasmid driving constitutive expression of HA-tagged anti-CD47 nanobody. **c**, pSC03 empty vector control.

**Supplementary Figure 2.**
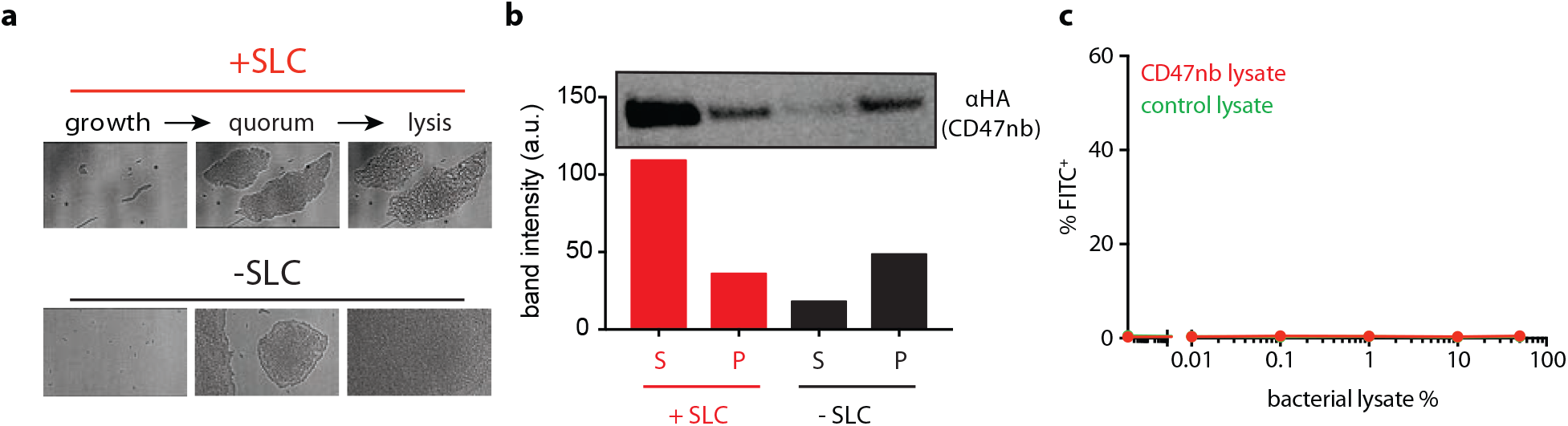
*E. coli* capable of synchronized lysis mediate production of blocking anti-CD47 nanobody. **a**, Bacterial growth dynamics over time in agar-pad microscope experiments. **b**, Immunoblot of bacterial culture supernatants (S) and cell pellets (P) in strains with and without SLC designed to constitutively produce HA-tagged CD47 nanobody. **c**, A20 cells were co-incubated with constant concentration of FITC conjugated IgG2a-FITC isotype control along with varying concentrations of bacterial lysates containing constitutively expressed CD47nb or empty vector.

**Supplementary Figure 3.**
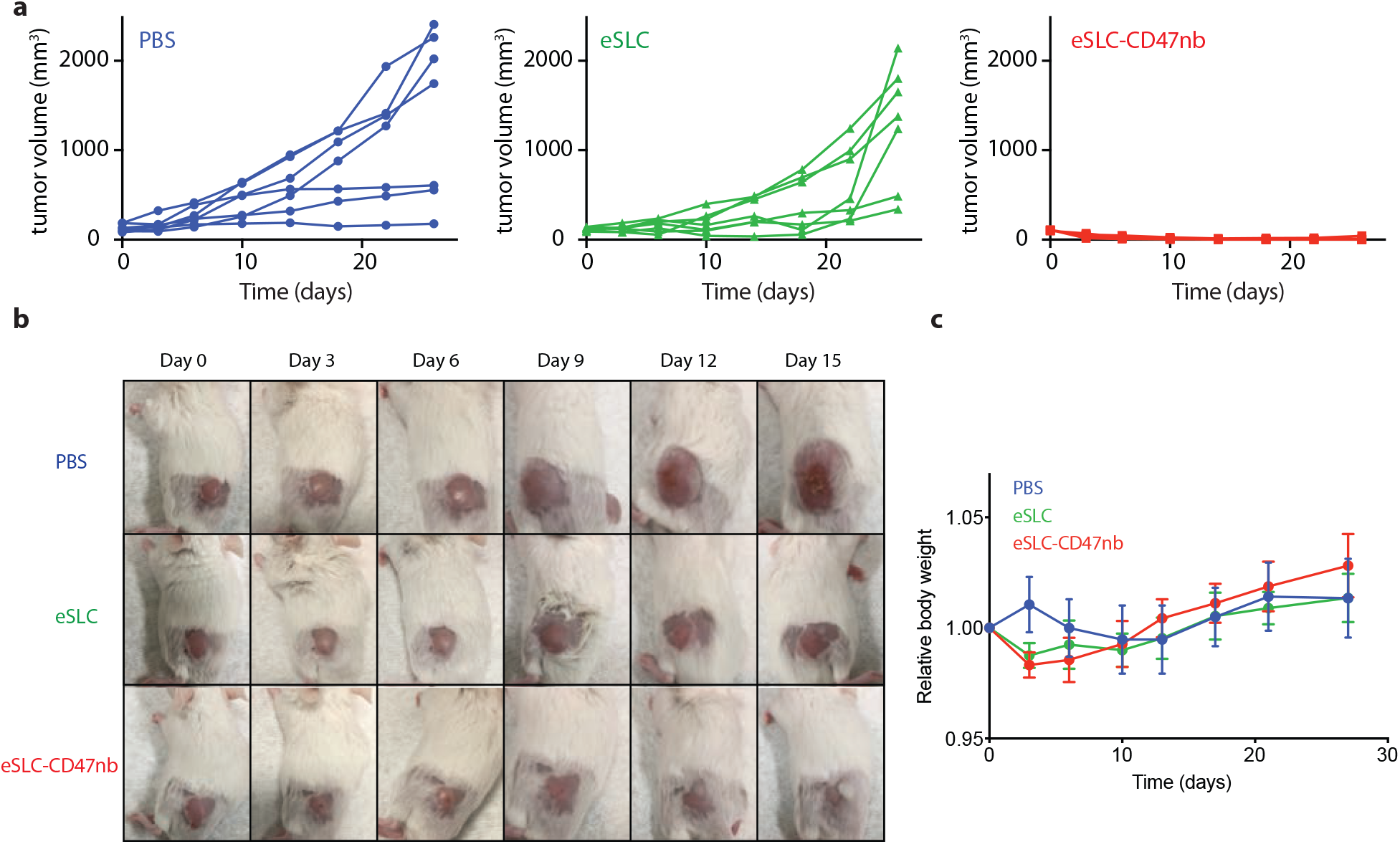
Parameters of intratumoral bacterial immunotherapy. **a**, Individual tumor growth trajectories. **b**, Representative images of subcutaneous A20 tumor bearing BALB/c mice treated with PBS, eSLC, or eSLC-CD47nb **c**, Relative body weight of A20 tumor bearing BALB/c mice over time.

**Supplementary Figure 4.**
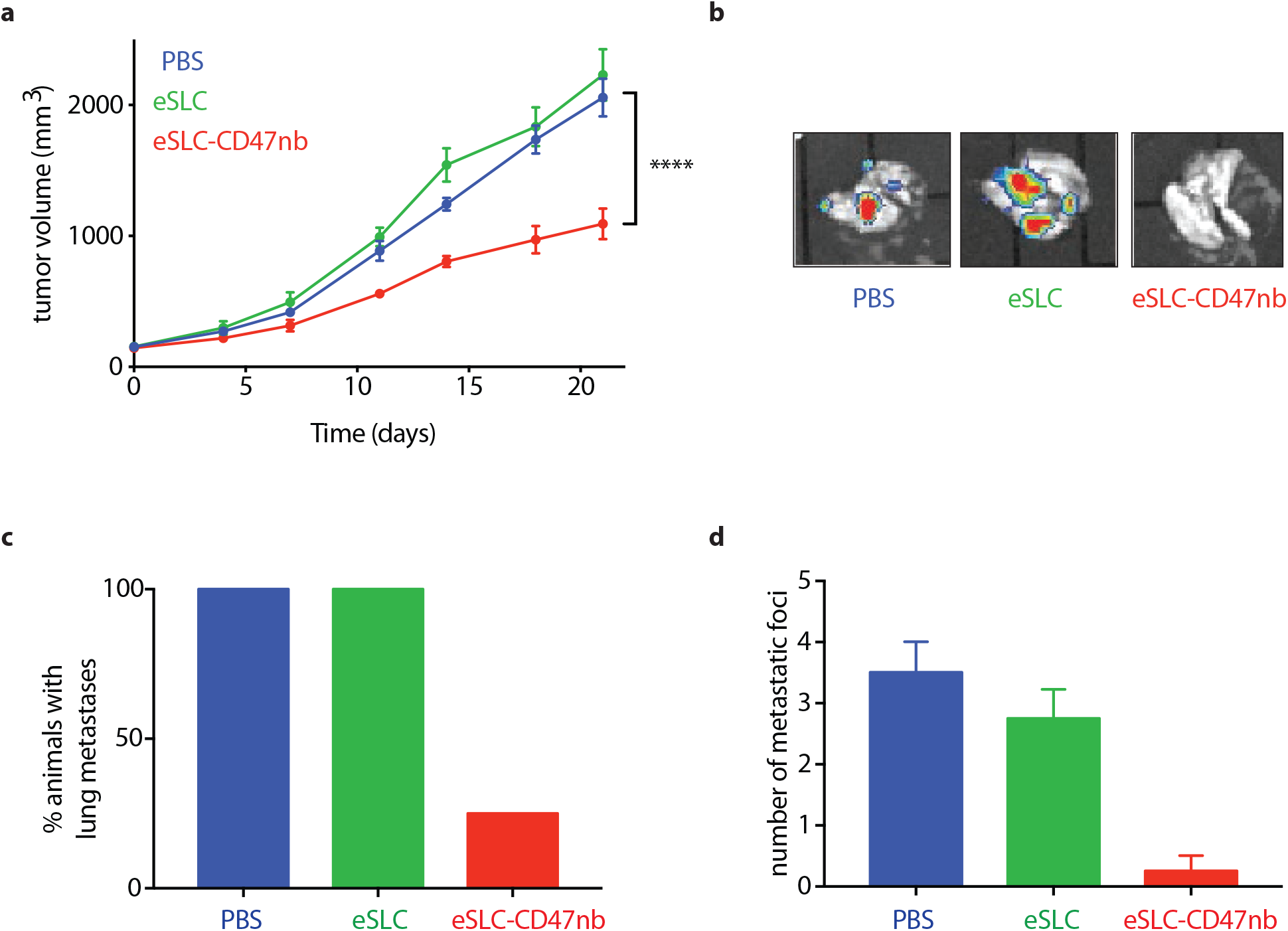
Immunotherapeutic bacteria limit tumor growth in a syngeneic murine model of triple negative breast cancer. **a**, 10^6^ 4T1-Luciferase mammary carcinoma cells were injected subcutaneously into the hind flank of 4-6-week-old BALB/c mice. When tumors were ~200 mm^3^ mice were randomized and received intratumoral injections of PBS, eSLC, or eSLC-CD47nb every 3 days for a total of 4 doses. Tumor growth curves. **b**, IVIS images of lungs extracted from mice bearing 4T1-Luciferase hind-flank tumors, **c**, Percentage of animals and **d**, number of 4T1-Luciferase metastatic foci in lungs of mice treated with PBS or eSLC or eSLC-CD47nb.

**Supplementary Figure 5.**
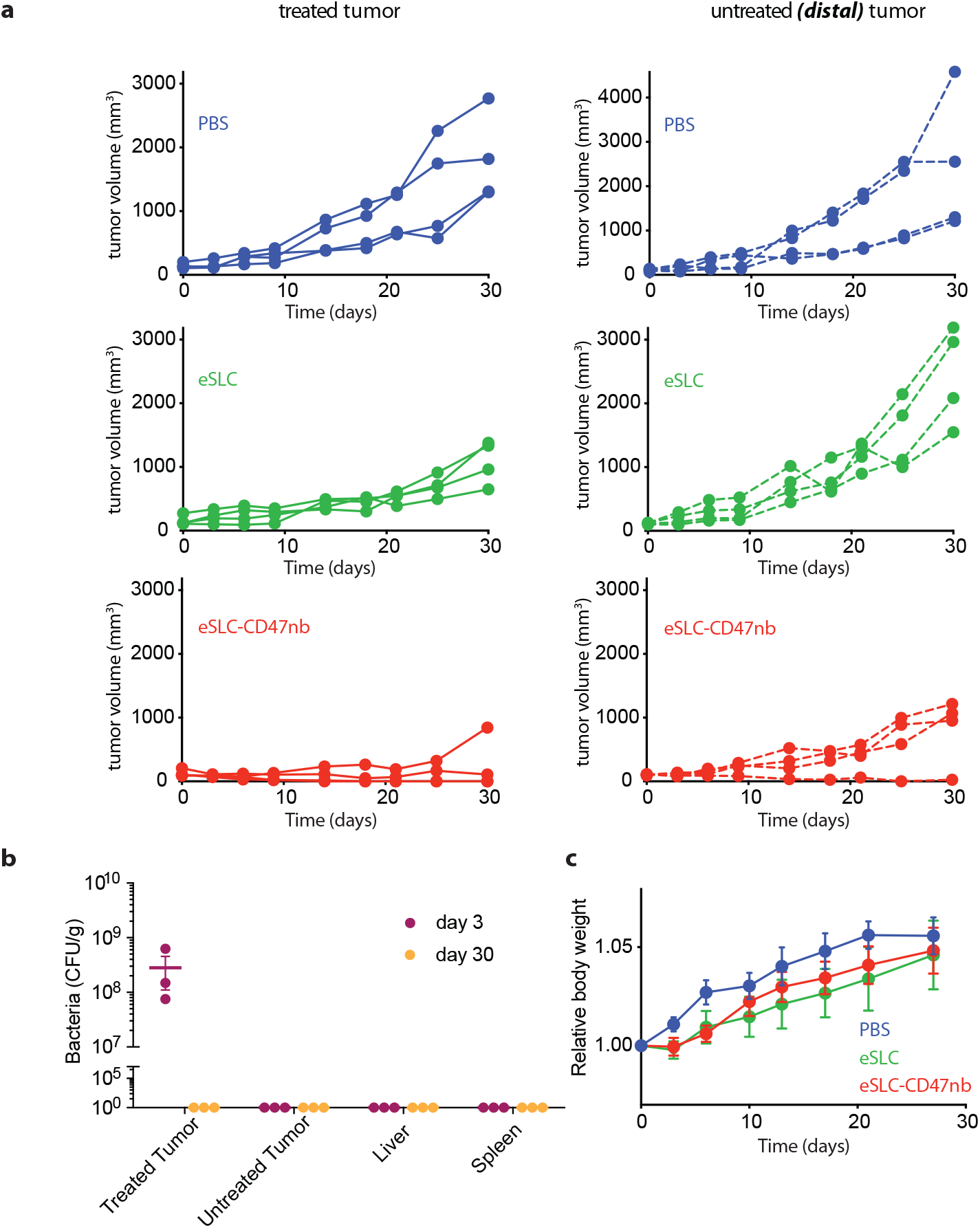
Intratumoral bacterial immunotherapy leads to distal tumor control. **a**, Individual tumor growth trajectories of treated (injected) and untreated A20 tumors following intratumoral PBS, eSLC, or eSLC-CD47nb injection. **b**, Biodistribution of eSLC-CD47nb *E. coli* on day 3 and day 30 following intratumoral bacterial injection. Excised tumors, livers and spleens were homogenized, serially diluted and plated on LB agar plates. Colonies were counted to determine CFU/g of tissue. **c**, Relative body weight of A20 tumor bearing BALB/c mice over time.

**Supplementary Figure 6.**
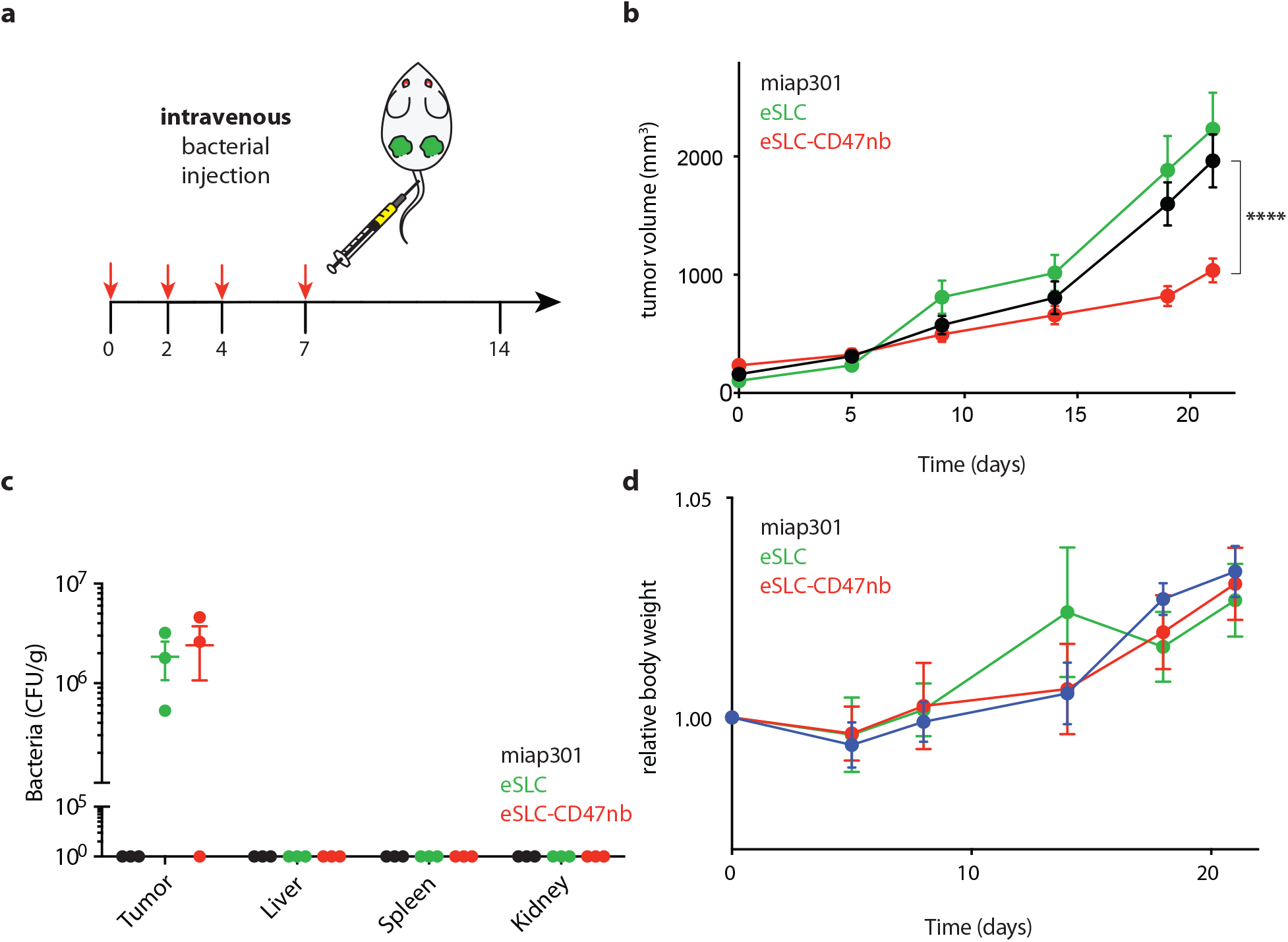
Intravenous bacterial immunotherapy limits tumor growth in a subcutaneous A20 lymphoma model. **a**, Treatment schedule. **b**, Mice were injected with 5 × 10^6^ A20 cells into both hind flanks. When tumor volume was 100mm^3^ – 200mm^3^ mice received intravenous injections of eSLC or eSLC-CD47nb or intraperitoneal injections of CD47mAb, miap301. **c**, Biodistribution of eSLC-CD47nb *E. coli* on day 8 following final intravenous bacterial treatment. Excised tumors, livers, spleens and kidneys were homogenized, serially diluted and plated on LB agar plates. Colonies were counted to determine CFU/g of tissue. **d**, relative body weight of A20 tumor bearing BALB/c mice receiving intravenous bacterial injections or intraperitoneal miap301 injections.

**Supplementary Figure 7.**
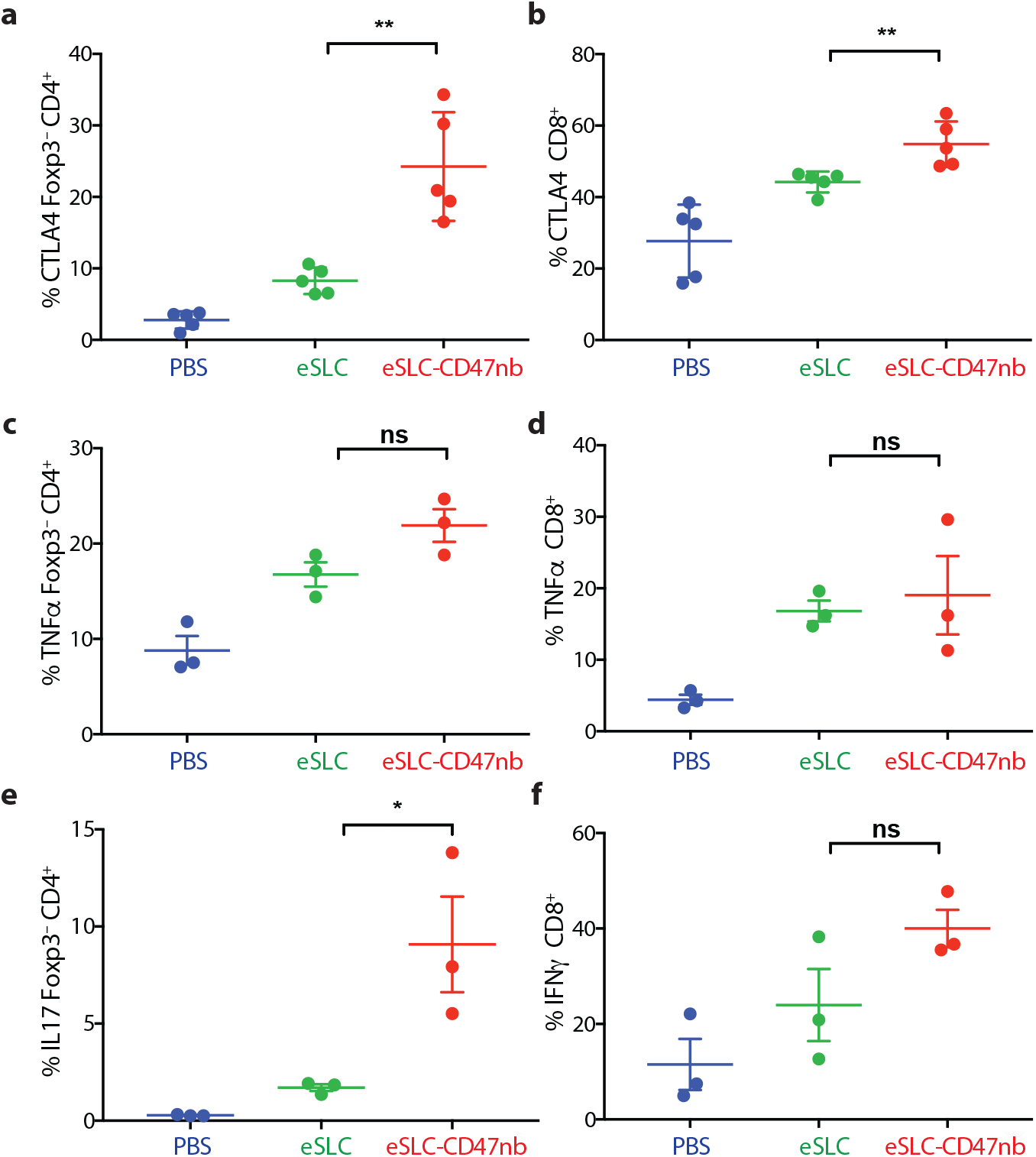
Immunophenotyping of tumor infiltrating lymphocytes following intratumoral bacterial injection. 5 × 10^6^ A20 cells were implanted into the hind flanks of BALB/c mice. When tumors reached ~100 mm^3^ in volume (day 0), mice were treated with either PBS, eSLC or eSLC-CD47nb on day 0, 4 and 7. Tumors were extracted and analyzed by flow cytometry on day 8. **a**, **b**, Percentages of CTLA4^+^ cells within Foxp3^−^CD4^+^ and CD8^+^ T cell respectively. **c**, **d**, Percentages of TNFα^+^ within Foxp3^−^ CD4^+^ and CD8^+^ T cells respectively following *ex vivo* stimulation. **e**, Percentage of IL17^+^ within Foxp3^−^CD4^+^ T cells following *ex vivo* stimulation. **f**, Percentages of intratumoral IFNγ^+^ within CD8^+^ T cells following *ex vivo* stimulation.

**Supplementary Figure 8.**
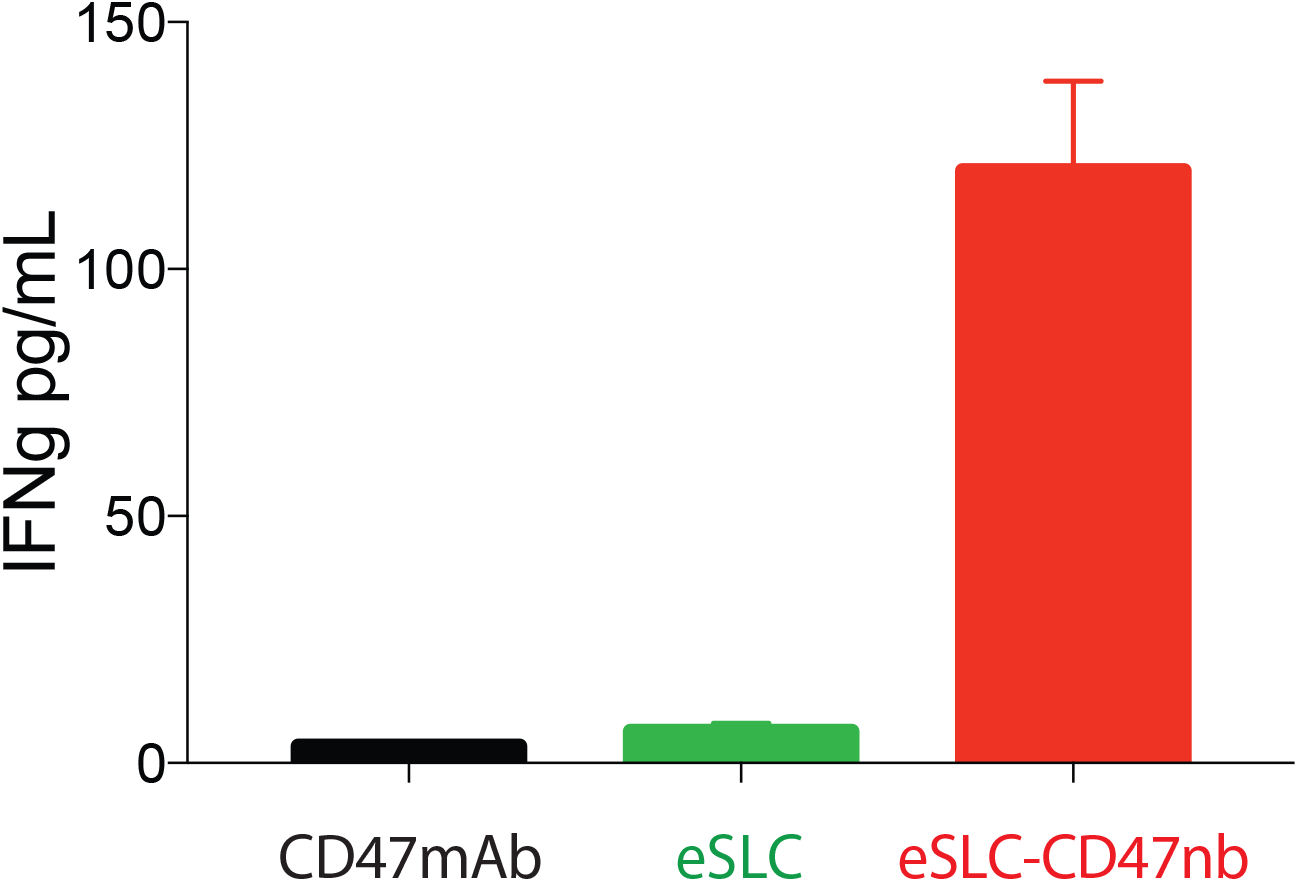
Immunotherapeutic bacteria leads to increased interferon-γ production by splenic T cells following stimulation with tumor antigens. IFN-γ ELISA of supernatants from overnight coincubation of isolated splenocytes with irradiated A20 cells.

**Supplementary Figure 9.**
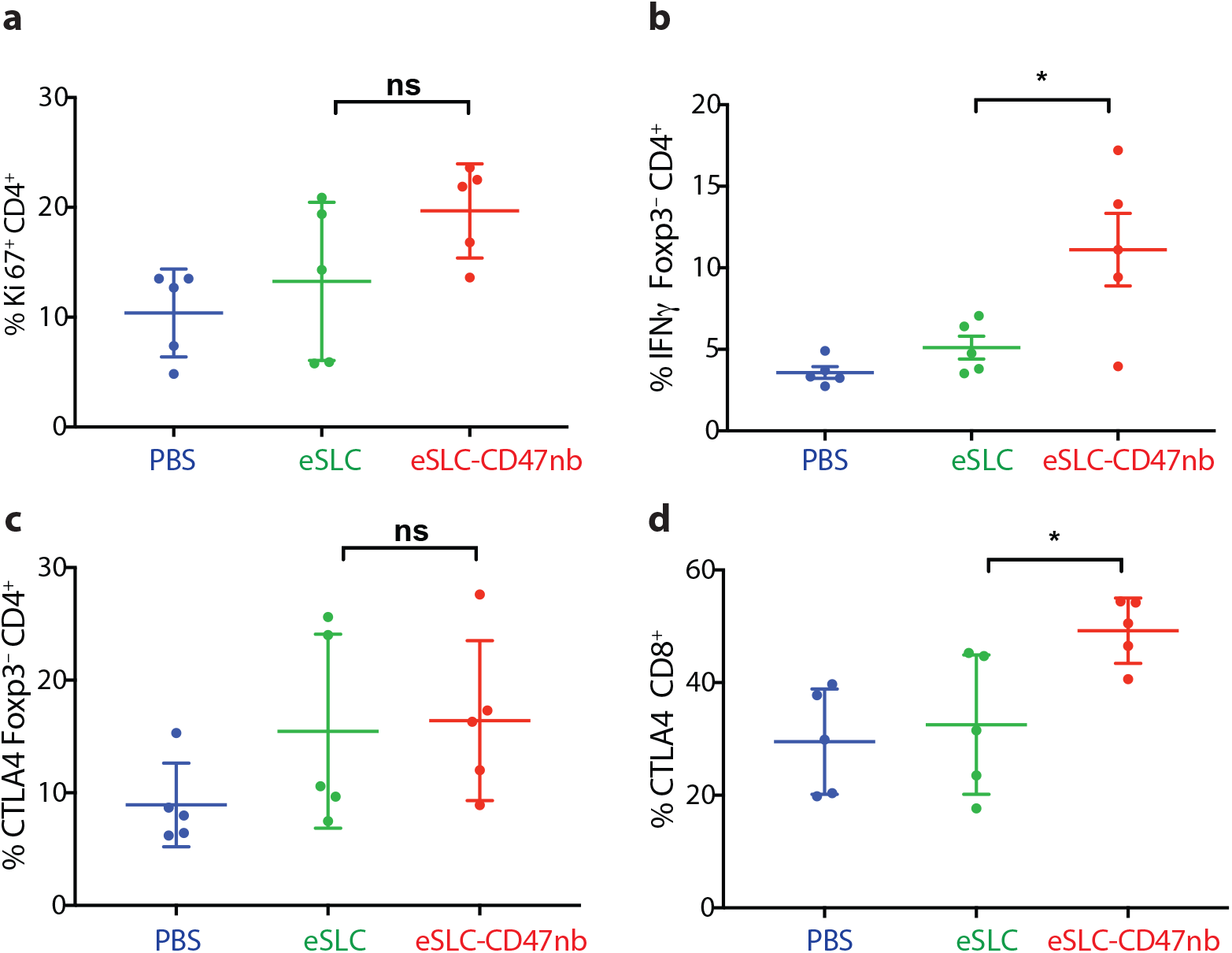
Immunophenotyping of tumor infiltrating lymphocytes in untreated tumors following single-flank bacterial injection. 5 × 10^6^ A20 cells were implanted into the hind flanks of BALB/c mice. When tumors reached ~100 mm^3^ in volume (day 0), mice were treated with either PBS, eSLC or eSLC-CD47nb on day 0, 4 and 7 into a single tumor. Untreated tumors were extracted and analyzed by flow cytometry on day 8. **a**, **b**, Frequency of Ki-67^+^ cell within Foxp3^−^CD4^+^ T cells. **b**, Frequency of tumor infiltrating IFNγ^+^ within Foxp3^−^CD4^+^ T cells following *ex vivo* stimulation with PMA and ionomycin in the presence of brefeldin A. **c**, **d**, Percentages of CTLA4^+^ within Foxp3^−^CD4^+^ T and CD8^+^ T cells compartments respectively.

## REFERENCES

1. Fischbach, M.A., Bluestone, J.A. & Lim, W.A. Cell-based therapeutics: the next pillar of medicine. Sci Transl Med 5, 179ps177 (2013).

2. Weber, W. & Fussenegger, M. Emerging biomedical applications of synthetic biology. Nat Rev Genet 13, 21–35 (2011).

3. Lim, W.A. & June, C.H. The Principles of Engineering Immune Cells to Treat Cancer. Cell 168, 724–740 (2017).

4. Ruder, W.C., Lu, T. & Collins, J.J. Synthetic biology moving into the clinic. Science 333, 1248–1252 (2011).

5. Chen, Y.Y. & Smolke, C.D. From DNA to targeted therapeutics: bringing synthetic biology to the clinic. Sci Transl Med 3, 106ps142 (2011).

6. Chien, T., Doshi, A. & Danino, T. Advances in bacterial cancer therapies using synthetic biology. Curr Opin Syst Biol 5, 1–8 (2017).

7. Din, M.O., et al. Synchronized cycles of bacterial lysis for in vivo delivery. Nature 536, 81-85 (2016).

8. Pedrolli, D.B., et al. Engineering Microbial Living Therapeutics: The Synthetic Biology Toolbox. Trends Biotechnol (2018).

9. Sockolosky, J.T., et al. Durable antitumor responses to CD47 blockade require adaptive immune stimulation. Proc Natl Acad Sci U S A 113, E2646–2654 (2016).

10. Majeti, R., et al. CD47 is an adverse prognostic factor and therapeutic antibody target on human acute myeloid leukemia stem cells. Cell 138, 286–299 (2009).

11. Willingham, S.B., et al. The CD47-signal regulatory protein alpha (SIRPa) interaction is a therapeutic target for human solid tumors. Proc Natl Acad Sci U S A 109, 6662–6667 (2012).

12. Coley, W.B. II. Contribution to the Knowledge of Sarcoma. Ann Surg 14, 199–220 (1891).

13. Berendt, M.J., North, R.J. & Kirstein, D.P. The immunological basis of endotoxin-induced tumor regression. Requirement for T-cell-mediated immunity. J Exp Med 148, 1550–1559 (1978).

14. Tsung, K. & Norton, J.A. Lessons from Coley’s Toxin. Surg Oncol 15, 25–28 (2006).

15. Mellman, I., Coukos, G. & Dranoff, G. Cancer immunotherapy comes of age. Nature 480, 480–489 (2011).

16. Jiang, S.N., et al. Inhibition of tumor growth and metastasis by a combination of Escherichia coli-mediated cytolytic therapy and radiotherapy. Mol Ther 18, 635–642 (2010).

17. Malmgren, R.A. & Flanigan, C.C. Localization of the vegetative form of Clostridium tetani in mouse tumors following intravenous spore administration. Cancer Res 15, 473–478 (1955).

18. Brown, J.M. & Wilson, W.R. Exploiting tumour hypoxia in cancer treatment. Nat Rev Cancer 4, 437–447 (2004).

19. Gardner, T.S., Cantor, C.R. & Collins, J.J. Construction of a genetic toggle switch in Escherichia coli. Nature 403, 339–342 (2000).

20. Basu, S., Gerchman, Y., Collins, C.H., Arnold, F.H. & Weiss, R. A synthetic multicellular system for programmed pattern formation. Nature 434, 1130–1134 (2005).

21. Friedland, A.E., et al. Synthetic gene networks that count. Science 324, 1199–1202 (2009).

22. Danino, T., Mondragon-Palomino, O., Tsimring, L. & Hasty, J. A synchronized quorum of genetic clocks. Nature 463, 326–330 (2010).

23. Elowitz, M.B. & Leibler, S. A synthetic oscillatory network of transcriptional regulators. Nature 403, 335–338 (2000).

24. Jaiswal, S., et al. CD47 is upregulated on circulating hematopoietic stem cells and leukemia cells to avoid phagocytosis. Cell 138, 271–285 (2009).

25. Liu, X., et al. CD47 blockade triggers T cell-mediated destruction of immunogenic tumors. Nat Med 21, 1209–1215 (2015).

26. Kauder, S.E., et al. ALX148 blocks CD47 and enhances innate and adaptive antitumor immunity with a favorable safety profile. PloS one 13, e0201832 (2018).

27. Liu, X., et al. Dual Targeting of Innate and Adaptive Checkpoints on Tumor Cells Limits Immune Evasion. Cell Rep 24, 2101–2111 (2018).

28. Huang, Y., Ma, Y., Gao, P. & Yao, Z. Targeting CD47: the achievements and concerns of current studies on cancer immunotherapy. J Thorac Dis 9, E168–E174 (2017).

29. Advani, R., et al. CD47 Blockade by Hu5F9-G4 and Rituximab in Non-Hodgkin’s Lymphoma. N Engl J Med 379, 1711–1721 (2018).

30. Ingram, J.R., et al. Localized CD47 blockade enhances immunotherapy for murine melanoma. Proc Natl Acad Sci U S A 114, 10184–10189 (2017).

31. Skinner, S.O., Sepulveda, L.A., Xu, H. & Golding, I. Measuring mRNA copy number in individual Escherichia coli cells using single-molecule fluorescent in situ hybridization. Nat Protoc 8, 1100–1113 (2013).

32. Haldimann, A. & Wanner, B.L. Conditional-replication, integration, excision, and retrieval plasmid-host systems for gene structure-function studies of bacteria. J Bacteriol 183, 6384–6393 (2001).

33. Gerdes, K., et al. Mechanism of postsegregational killing by the hok gene product of the parB system of plasmid R1 and its homology with the relF gene product of the E. coli relB operon. EMBO J 5, 2023–2029 (1986).

34. Derman, A.I., et al. Alp7R regulates expression of the actin-like protein Alp7A in Bacillus subtilis. J Bacteriol 194, 2715–2724 (2012).

